# Systemic infections alter cortical transcriptionalsignatures in Alzheimer’s disease

**DOI:** 10.1101/2025.10.23.682825

**Authors:** Giulia Pegoraro, Lachlan F. MacBean, Adam R. Smith, Darren M. Soanes, Rebecca G. Smith, Jennifer Imm, Joshua Harvey, Morteza Kouhsar, Valentin Laroche, Luke Weymouth, Isabel Castanho, Winston Hide, Laura Palmer, Debra J. Lett, Andrew C. Robinson, Natalie Woodman, Karina McDade, Delphine Boche, Ehsan Pishva, Katie Lunnon

**Author notes:** Corresponding author: Katie Lunnon, University of Exeter, RILD, Barrack Road, Exeter, Devon, UK. EX2 5DW.

## Abstract

Alzheimer’s disease (AD) is characterized by neuroinflammation, yet the impact of concurrent systemic infections on the AD brain remains poorly understood. We investigated the molecular mechanisms underlying the central nervous system response to systemic infections in AD by analyzing RNA sequencing data generated in the prefrontal cortex from 202 post-mortem donors (113 AD, 89 controls), where we stratified by the presence of a respiratory infection at the time of death. We identified 763 significantly differentially expressed genes (DEGs) between AD and controls without infection, which were enriched for oxidative phosphorylation and neurodegenerative pathways. In contrast, 122 DEGs distinguished AD from controls during infection, with 57 genes uniquely altered in AD in the presence of infection, including *MAPK4*, *VAV3*, and *POU3F4*, implicating infection-dependent mechanisms of vascular and immune regulation. Pathway activity analysis revealed that infection in AD suppresses some immune and vascular pathways, while enhancing transcriptional and developmental programs. Weighted gene co-expression network analysis uncovered three key modules: one module strongly associated with AD, enriched for aging and signal transduction; one module linked to both AD and infection, highlighting cytoskeletal remodelling and host–pathogen interactions; and one module specific to infection, enriched in astrocytes, pericytes, and endothelial cells, implicating blood–brain barrier dysfunction. These findings suggest that systemic respiratory infections reshape transcriptional programs in the AD brain, dampening immune effector pathways and engaging vascular and host–pathogen processes in blood–brain-barrier–associated cell types. Our results highlight the complex interplay between systemic infection, neuroinflammation, and vascular responses in AD.

## Introduction

Alzheimer’s Disease (AD) is a progressive neurodegenerative disorder that accounts for over 60-70% of dementia cases globally (World Health Organization, 2017). The pathological hallmarks of AD include the deposition of beta-amyloid (Aβ) plaques, neurofibrillary tangle (NFT) formation, and neuroinflammation, all of which contribute to synaptic dysfunction, neuronal loss, and cognitive decline (Akiyama et al., 2000; Mandelkow and Mandelkow, 1998; Murphy and LeVine, 2010). Neuroinflammation, driven primarily by the activation of microglia and astrocytes within the central nervous system (CNS), plays a multifaceted role in AD pathogenesis. Although microglia play a key protective role in Aβ clearance, chronic neuroinflammation, characterized by a sustained release of pro-inflammatory cytokines, can exacerbate neuronal damage, ultimately accelerating disease progression (Kinney et al., 2018; Onyango et al., 2021). The innate immune response in AD is not limited to the brain; subtle alterations in the abundance of specific white blood cell types have been reported (Lunnon et al., 2012). Furthermore, plasma levels of some pro-(IL6, TNFα) and anti-inflammatory (IL1ra, IL10, IL13) cytokines have been inversely correlated with the level of brain atrophy in AD (Leung et al., 2013).

Recent evidence suggests a link between systemic inflammation and neuroinflammation in AD (Bayraktaroglu et al., 2025). Systemic inflammation, often triggered by peripheral infections, is characterized by the increased production of pro-inflammatory cytokines in the blood. These cytokines are hypothesized to access the CNS through a compromised blood-brain barrier (BBB), which becomes more vulnerable with aging and in neurodegenerative diseases (Galea, 2021; Zenaro et al., 2017). Once in the CNS, these circulating cytokines can further activate microglia and other glia, further exacerbating a pro-inflammatory environment, which in turn could contribute to neuronal dysfunction and cognitive decline (Doroszkiewicz et al., 2022).

A growing body of research has explored the relationship between infections and AD, typically focusing on specific pathogens. Pathogens such as *Herpes Simplex Virus 1* (HSV-1), *Epstein-Barr Virus* (EBV), and *Chlamydia pneumoniae* have been implicated in increasing the risk of AD development (Al-Atrache et al., 2019; Carbone et al., 2014; Harris and Harris, 2018). The COVID-19 pandemic has further highlighted the potential role of viral infections in neurodegeneration. Studies of *SARS-CoV-2* have demonstrated its capacity to invade the CNS via infected lymphocytes and olfactory neurons, mechanisms also observed in *Chlamydia pneumoniae* infections (Boelen et al., 2007; Hascup and Hascup, 2020). Additionally, *SARS-CoV-2* has been linked to an increased risk of a diagnosis of AD, with a higher risk in individuals over 85 years old (Wang et al., 2022) and increased cognitive impairment in individuals with dementia (Dubey et al., 2023). Clinical studies further support an association between systemic infections and AD manifestation and progression, with increased serum TNFα and IL1β and an increased rate of cognitive decline observed for several months following infection in individuals with AD (C et al., 2009, n.d.). Moreover, respiratory infections are a common cause of mortality in people with AD, with 46% of dementia-related deaths attributed to respiratory infections compared to 7% in age-matched non-demented individuals (Brunnström and Englund, 2009). Interestingly, several retrospective studies have highlighted that vaccination against common infections may reduce the risk of developing AD (Harris et al., 2023; Ukraintseva et al., 2024; Verreault et al., 2001).

Despite mounting evidence connecting systemic inflammation to neuroinflammation and cognitive decline in AD, the mechanisms underlying this are incompletely understood. Although numerous have systematically assessed the brain transcriptome in AD (Mathys et al., 2019; Mostafavi et al., 2018; Q et al., 2021; Raj et al., 2018; Wan et al., 2020), none of these studies have explored the impact of a concurrent systemic infection on these signatures, which is important given that clinical evidence suggests systemic infections can drive neuroinflammation and disease processes. Therefore, in the current study we have investigated the impact of systemic infections on gene expression signatures in AD brain by analysing 202 post-mortem prefrontal cortex samples from late-stage AD patients and control subjects, where we stratified by the presence of a respiratory infection at the time of death. This has allowed us to identify the molecular mechanisms specifically altered in AD brain during systemic infections.

## Methods and Materials

### Sample collection

Our study utilized post-mortem brain samples collated from five brain banks in the UK Brain Bank Network (UKBBN): the Edinburgh Brain Bank, the Manchester Brain Bank, the Newcastle Brain Tissue Resource, the Queen Square Brain Bank and the South West Dementia Brain Bank. Our cohort consisted of four groups divided as follows: (1) AD donors with an infection at the time of death, (2) AD donors without an infection at the time of death, (3) control donors with an infection at the time of death, and (4) control donors without an infection at the time of death. All AD donors had a pathological diagnosis of AD, with a Braak NFT stage of V-VI, whilst control donors had a Braak NFT stage ≤II. We identified donors with infections based on the clinical reports at the time of death, which specified the type of infection; any case in which infection was listed as the cause of death was classified as an infection case. Non-respiratory infections were excluded.

We excluded individuals with other types of dementia or other brain-related conditions, such as schizophrenia or brain injury. All donors included in the study were > 60 years to exclude early-onset AD, and groups were matched for age and sex as closely as possible.

### Sample preparation and nucleic acid extraction

The prefrontal cortex tissue was cut to collect 30mg on dry ice, before being milled over liquid nitrogen. The Qiagen AllPrep DNA/RNA/miRNA Universal kit was used to isolate DNA and total RNA from the dissected tissue according to the manufacturer’s instructions with minor modifications, namely that RNA was eluted by the addition of 2 × 25μL of RNase-free water. RNA was quantified and quality control (QC) checked using the Agilent Tapestation 4200 RNA screen tape, to assess RNA concentration and RNA integrity number (RIN). Two samples failed the QC step and were therefore removed. RNA was then stored at −80°C until required. All experiments were performed with the group blind to the user.

### RNA sequencing and data processing

The library preparation for the RNA sequencing (RNA-seq) was performed using the Illumina Stranded Total RNA Prep protocol using 9ng total RNA, with samples organised across plates by RIN score, whilst randomizing for other variables. The sequencing was performed in two batches using the Illumina NovaSeq System 6000 with 50 bp paired-end reads with ∼20 million reads per sample.

All sequencing data processing was conducted on a Unix-based operating system server at the University of Exeter. The FASTQ files containing the sequenced data underwent QC analysis using *FastQC* (“FastQC,” 2015). Subsequently, *FastQ* Screen was utilized to assess typical contaminants, and *fastp* was employed for trimming adapters and removing low-quality reads (Chen et al., 2018; Wingett and Andrews, 2018). Transcript expression quantification was performed using *Salmon* (version 1.8) (Patro et al., 2017), resulting in a gene count matrix. Instead of using RIN for analysis, we opted to utilize the transcript integrity number (TIN), which evaluates RNA integrity at the transcript level, addressing some limitations associated with RIN, such as its limited sensitivity for measuring degraded RNA (Wang et al., 2016). TIN was calculated using the tin.py function in the Python package *RSeQC* (version 5.0.0) (Wang et al., 2012). R (version 4.5) and Bioconductor (version 3.21) were utilized for all subsequent statistical analyses.

The genes in the count matrix underwent expression level filtration using the *filterByExpr* function in *EdgeR* (version 4.6.3), with a minimum count set to 10 and the minimum proportion of samples where a gene needs to be expressed set to 80%. This resulted in a final set of 16,593 genes. Data were normalized by taking the log2 of the values obtained from the counts per million (CPM) method. Normalization factors were computed based on library sizes using the trimmed mean of the M-values method (TMM) with the *calcNormFactors* function. Subsequently, outliers were identified using the Mahalanobis distance function. Principal component analysis (PCA) was conducted on the normalized expression data. A correlation matrix containing the correlation between the principal components (PCs) and the phenotypic and technical data associated with the samples was generated. The correlation matrix revealed strong correlations between the first PC and library size, TIN score, and sequencing plate, and so *ComBat* was used in the *SVA* library (version 3.56.0) to remove these batch effects prior to differential expression analysis.

### Differential expression analysis

Following data QC and after restricting our analysis to exclude donors who had died from a non-respiratory infection (to reduce sample heterogeneity), we were left with a final cohort of 202 individuals (Table 1). The package *countToFKPM* (version 1.04) was used to normalize the data to fragments per kilobase million (FKPM) (Alhendi, 2019), with CIBERSORT (version 1.04) (Newman et al., 2015) used to quantify the following cell types: neurons, microglia, oligodendrocyte precursor cells (OPC), oligodendrocytes, astrocytes, and endothelial cells (Yu and He, 2017). We used *limma* (version 3.64.3) (Ritchie et al., 2015) for differential expression analysis, fitting per-gene linear models with AD status, infection status, and their interaction as primary predictors, and including brain bank, age, sex, and estimated cell-type proportions as covariates. After fitting the model, comparisons were made between the following groups to identify differentially expressed genes (DEGs): AD donors with versus without an infection; control donors with versus without an infection; AD versus control donors with an infection; and AD versus control donors without an infection. P-values were adjusted for multiple testing using the Benjamini-Hochberg (BH) method, a false discovery rate (FDR)-controlling method, with these values referred to herein as p_adj_.

**Table 1:** Demographic information for the cohort stratified by diagnostic group. . For each group, we provide the total number of Control and Alzheimer’s disease (AD) samples used in the analysis, mean age at death (± standard deviation (SD), years), sex distribution (Female (F) / Male (M)), mean transcript integrity number (TIN ± SD), mean post-mortem interval (PMI ± SD, hours).

| Characteristic | No systemic infection |  | With systemic infection |  |
| --- | --- | --- | --- | --- |
|  | Control | AD | Control | AD |
| Sample number | 52 | 55 | 37 | 58 |
| Age | 80 ± 9 | 81 ± 7 | 87 ± 9 | 82 ± 8 |
| Sex (F/M) | 23 / 29 | 30 / 25 | 17 / 20 | 35 / 23 |
| TIN score | 62.6 ± 4.5 | 61.5 ± 4.7 | 64.1 ± 3.7 | 62.7 ± 4.6 |
| PMI | 45 ± 21 | 41 ± 26 | 57 ± 37 | 47 ± 25 |

### Co-expression network analysis

Weighted gene co-expression network analysis (WGCNA) was performed using the R package *WGCNA* (version 1.73) to identify co-expressed modules and then relate these to disease and infection status (Langfelder and Horvath, 2008). Before the analysis, we regressed out the same covariates as in our differential gene expression analysis (*i.e.,* age, sex, cell proportions, and brain bank). We selected a soft threshold of five using the function *pickSoftThreshold* within the *WGCNA* package. Following the calculation of the adjacency matrix, it was converted into a topological overlap matrix (TOM). Modules were subsequently formed using hierarchical clustering with the following parameters: a minimum module size (*minModuleSize*) of 50, a maximum block size (*maxBlockSize*) of 20,000 and a signed network type. Each module was assigned an arbitrarily chosen colour, with the grey module excluded as it contains all unassigned genes. Module eigengenes (MEs) were calculated for each module and represented the overall expression behavior of the genes within the module. The co-expression modules were then associated with AD and infection status using pairwise Pearson correlation analysis. For modules that contained >1,000 genes, we utilized “hub” genes for downstream analysis. Hub genes were defined as those within a module that had a module membership (MM) ≥ the 50th percentile and gene significance (GS) ≥ the 75th percentile for the trait of interest. In the case of modules that showed an association with both AD and infection status, GS was computed separately for each trait, and then we took forward genes that showed a GS ≥ the 75th percentile for both traits and a MM ≥ the 50th percentile.

### Downstream pathway, cell-type enrichment and functional module analysis

We performed pathway enrichment analysis utilizing the *clusterProfiler* package (version 4.16.0) (Wu et al., 2021). Specifically, an overrepresentation analysis (ORA) was conducted using Gene Ontology (GO) Biological Processes terms and Kyoto Encyclopedia of Genes and Genomes (KEGG), with the background gene set defined as the 16,593 genes that passed QC. Multiple testing correction was performed using the FDR adjustment to yield q-values.

To assess the direction of changes in biological pathways, we performed Gene set variation analysis (GSVA) to calculate the pathway activity across our cohort, in combination with linear modelling with *limma*. GO Biological Processes served as the reference database for pathway definitions. GSVA was performed using the *GSVA* R package (version 2.2.0) and was applied to estimate sample-wise enrichment scores for these pathways, using the Gaussian kernel method (Hänzelmann et al., 2013). Differential activity of the identified pathways was tested using *limma* using the same model design and comparisons used for the differential gene expression analysis.

To determine cell type enrichment for genes annotated to modules of interest, we performed expression-weighted cell-type enrichment analysis (EWCE) (Skene and Grant, 2016). The single-cell type transcriptome reference datasets utilized in this analysis were obtained from previous studies of human prefrontal cortex (Mathys et al., 2019). The reference datasets were first prepared using the *drop_uninformative_genes()* function, which removes uninformative genes to reduce computation time and noise. Following this filtering, the single-nucleus RNA-seq (snRNA-seq) datasets used in EWCE (version 1.16.0) were generated by the *generate_cell_type_data()* function. Cell-type enrichment analysis was then performed using the *bootstrap_enrichment_test()* function, with 100,000 bootstrapping repetitions. Significant cell types were defined as those with a p_adj_ < 0.05 after BH correction.

In our WGCNA analysis, to identify key functional sub-modules relevant to AD, we applied a functional module discovery method contained in the HumanBase platform (Krishnan et al., 2016) to WGCNA modules of interest. This approach identifies clusters of genes exhibiting coordinated activity, annotated with biological processes from GO. As HumanBase allows the analysis of tissue-specific networks, we selected the cerebral cortex as the tissue of interest, given that we had used prefrontal cortex samples in our study.

### Data and code availability

RNA-seq will be made available on the Gene Expression Omnibus (GEO) Platform upon final publication. Analysis scripts used in this manuscript are available at https://github.com/UoE-Dementia-Genomics/ADInfection_transcriptomics_analysis.

## Results

### Systemic infection in AD induces distinct transcriptional signatures in the brain

To investigate whether systemic infection alters molecular pathways in AD brain we conducted differential expression analyses comparing AD cases to controls, stratifying by the presence of a respiratory infection at the time of death. First, we explored gene expression alterations in AD brain compared to controls in the absence of systemic infection, to identify the molecular mechanisms specifically associated with AD. We identified 763 significant DEGs (p_adj_ < 0.05) in AD cases compared to controls in the absence of infection (Supplementary Table 1, Fig. 1A), with the ten most significant genes including *ADAMTS2*, *HEY1*, *CIC*, *VGF*, *ZNF704*, *LINC01322*, *PPDPF*, *KBTBD12*, and two unannotated transcripts (ENSG00000254746 and ENSG00000260293). Pathway enrichment analysis on these 763 genes showed a significant enrichment in three GO biological processes pathways, all related to energy metabolism (oxidative phosphorylation: q = 0.0233, aerobic respiration: q = 0.0246, generation of precursor metabolites and energy: q = 0.0246, Supplementary Table 2), whilst we identified four significant KEGG pathways (oxidative phosphorylation: q = 1.87 x 10^-3^, diabetic cardiomyopathy: q = 0.0228, Huntington’s disease (HD): q = 0.0228, prion disease: q = 0.0228, Supplementary Table 3). This enrichment for HD and prion disease likely represents an enrichment for neurodegenerative disease pathways, as we also saw a nominally significant enrichment for AD (p = 0.0177), which did not reach FDR significance, but that featured a large number of overlapping genes with those the HD and prion disease pathways. To explore whether specific biological pathways are dysregulated in AD when there is no concurrent systemic infection we performed GSVA using GO Biological Process terms to quantify pathway activity per sample and then compare these between AD cases and controls without a respiratory infection at the time of death. This pathway centric analysis identified nine pathways that showed a significant increase in activity, including B cell receptor signalling and signal transduction in response to DNA damage (Fig. 1C). Twenty pathways showed down-regulation of activity, included those related to immune cell functions, such as phagosome-lysosome fusion and antimicrobial humoral immune response mediated by antimicrobial peptide (p_adj_ < 0.05, Fig. 1C).

**Fig. 1:**
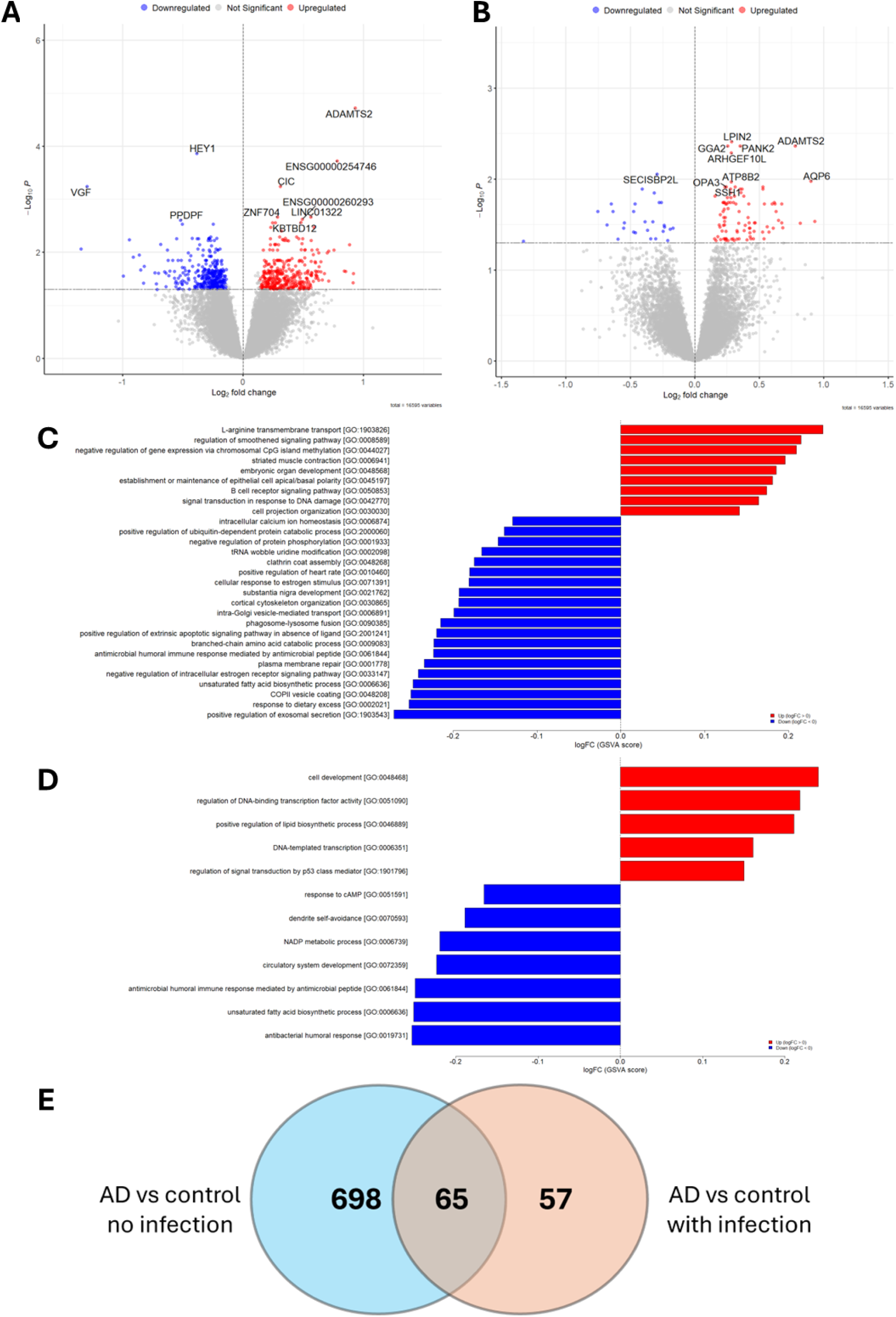
Differentially expressed genes in AD cases compared to controls stratified by the presence of a respiratory infection at the time of death. Volcano plots showing differentially expressed genes in AD cases compared to controls, in (A) the absence, and (B) the presence of a respiratory infection at the time of death. False discovery rate (FDR) significant genes (p_adj_ < 0.05) are shown in blue (down-regulated) and red (up-regulated) above the horizontal dashed line. The ten most significant genes are labelled with gene name or, for non-coding genes, the ensembl ID. Pathways identified in Gene Set Variation Analysis (GSVA) exploring pathway activity in AD compared to control in (C) the absence, and (D) the presence of a respiratory infection at the time of death. Blue bars represent pathways showing an FDR-significant down-regulation of activity, whilst red bars represent an FDR significant up-regulation of activity. (E) A Venn diagram showing the number of FDR-significant differentially expressed genes identified and overlapping between the different comparisons.

When we compared DEGs in AD brain compared to controls, where there was a concurrent systemic infection at the time of death we identified 122 significant genes (Supplementary Table 4, Fig. 1B), with the ten most significant genes corresponded to *LPIN2, ADAMTS2, GGA2, PANK2, ARHGEF10L, SECISBP2L, AQP6, ATP8B2, SSH1 and OPA3*. We did not identify any FDR-significant GO (Supplementary Table 5) or KEGG pathways (Supplementary Table 6). However, using GSVA to explore pathway activity, we did identify five up-regulated pathways, including those related to transcription and development, and seven down-regulated pathways, including those related to immune cell responses and vascular development, such as humoral responses and circulatory system development (p_adj_ < 0.05, Fig. 1D).

Next, we wanted to identify the distinct transcriptomic signatures that are observed in AD brain in the presence of a systemic infection but do not occur in the absence of an infection. Notably, 57 genes were differentially expressed when comparing AD and control samples with an infection and were not differentially expressed (p_adj_ > 0.05) in AD the absence of infection (Supplementary Table 7, Fig. 1E). Although 39 of these genes showed a nominally significant change in the absence of infection, 18 showed no nominal transcriptional change (p < 0.05), suggesting these are infection-dependent transcriptional changes in AD. This included *OPA3*, *VAV3*, *SNX21*, *KIFC3*, *CBS*, *AP5S1*, *MARCHF1*, *SP8*, *TNR*, *POU3F4*, *TMLHE*, *RFLNB*, *TATDN2*, *RIMKLB*, *MAPK4*, *CENPP* and two unannotated transcripts (ENSG00000279168 and ENSG00000271793). Despite identifying these infection-dependent differentially expressed genes in AD, we did not identify any FDR-significant differentially expressed genes when comparing AD cases with infection to AD cases without infection (Supplementary Table 8), nor when comparing infected controls to uninfected controls (Supplementary Table 9), suggesting that the 57 genes we identified in AD during systemic infection are driven by the influence of both AD and infection.

### WGCNA reveals co-expression modules linked to AD and infection

Next, we used WGCNA to identify modules of co-expressed genes associated with disease status and/or the presence of infection, with the rationale being that this may reveal important functional pathways altered in the brain in AD, and during systemic respiratory infections. We identified 12 modules of co-expressed genes (Fig. 2A), with one showing a Bonferroni-significant correlation with AD status (red, r = 0.28, p = 4 x 10^-5^). This module consisted of 568 genes and although pathway enrichment analysis showed no significant GO pathways (Supplementary Table 10), we did identify 25 significant KEGG pathways, many related to aging, signal transduction, and endocrine systems (Supplementary Table 11, Fig. 2B).

**Fig. 2:**
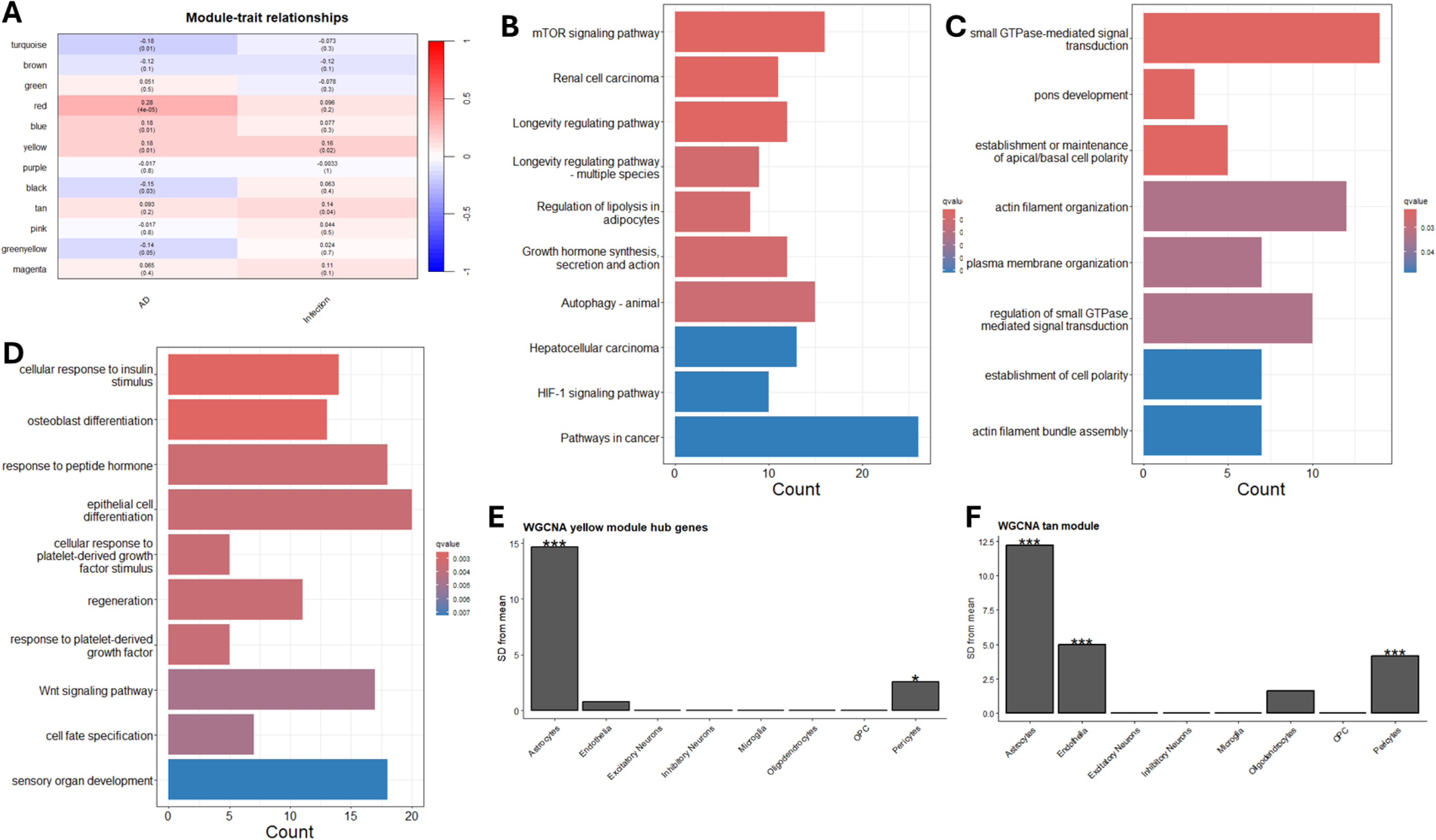
Modules of co-expressed genes associated with AD and infection. (**A**) Weighted gene correlation network analysis (WGCNA) module–trait matrix showing Pearson correlations (r, upper number) with unadjusted p-value (lower number in parentheses). (**B**) The ten most significant KEGG pathways enriched in the red module (of the 25 that passed the FDR significance threshold). Bar length indicates the number of genes annotated to each term; fill colour reflects adjusted p-values. (**C**) The eight FDR significant GO Biological Process pathways enriched in yellow-module hub genes. (**D**) The ten most significant GO Biological Process terms enriched in the tan module (of the 78 that passed the FDR significance threshold). Expression weighted cell-type enrichment (EWCE) results of (**E**) yellow-module hub genes and (**F**) tan module genes. Bars show standard deviation (SD) from the mean; asterisks denote significance, demonstrating which cell types are enriched (*p_adj_ < 0.05, **p_adj_ < 0.01, ***p_adj_ < 0.001).

As no studies to date have explored transcriptomic signatures in the human brain associated with systemic infection, despite it having epidemiological links to AD, we were also interested in exploring modules associated with infection status. Although we did not identify any Bonferroni-significant correlations for the 12 modules with infection status, we did identify two nominally significant modules (yellow: r = 0.16, p = 0.02; tan: r = 0.14, p = 0.04, Fig. 2A). We were particularly interested in the yellow module, consisting of 1,479 genes, as this also as this also showed a nominally significant association with AD status (r = 0.18, p = 0.01). Interestingly, this module contained thirty genes that overlapped with the 122 genes we had identified in our differential gene expression analysis comparing AD and control subjects with infection. This included the genes *LAMP1, RBM5, ATF2, PAK4, PRKCH, VAMP3 and ACTN4*. As the module was large (> 1,000 genes) we extracted the hub genes (N = 124) for downstream analysis as these represent highly connected genes that pay a critical role in the module. These were defined as the genes showing a MM > 0.5 and PS > 0.75 for both AD and infection status). GO enrichment analysis highlighted eight significant pathways, including several related to signal transduction and cellular organization (Supplementary Table 12, Fig. 2C), with no significant KEGG terms (Supplementary Table 13). When we performed pathway enrichment analysis on the 188 genes in the tan module we identified 78 significant GO terms (Supplementary Table 14, Fig. 2D), but no significant KEGG terms (Supplementary Table 15), with many of these relating to signalling pathways and cellular responses.

### Glial and neurovascular cells show transcriptomic alterations during systemic infection

To identify the cell types underlying the transcriptomic patterns in the three modules where we had seen a correlation with AD or infection (red, yellow, tan), we performed EWCE using snRNA-seq data previously generated in human prefrontal cortex [38]. For the yellow module that was nominally significantly associated with both AD and systemic infection, we observed a significant enrichment of hub genes in astrocytes and pericytes (Fig. 2E), whilst for the tan module, which was associated with infection only, genes were enriched in astrocytes, pericytes and endothelial cells (Fig. 2F). In contrast, genes in the red module, which showed a Bonferroni significant correlation with AD status but not infection, were not enriched for any particular cell type, suggesting broad cellular changes.

### Functional sub-module analysis reveals specific pathways associated with AD and infection

To further characterize the yellow, tan and red modules we performed functional module detection to identify “sub-modules” associated with specific pathways within each WGCNA module. In the yellow module, which had been associated with both AD and infection, we identified ten sub-modules (M1A–M10A) (Fig. 3). Unsurprisingly, immune and inflammatory signalling pathways featured prominently: M4A captured I-κB kinase/NF-κB and tumour necrosis factor responses; M2A centred on mast cell activation/degranulation; and M1A encompassed receptor transactivation pathways, including epidermal growth factor receptor (EGFR) transactivation by G-protein coupled receptors (GPCRs) and ciliary neurotrophic factor (CNTF)-mediated signalling. Two sub-modules mapped to RNA regulatory programmes—M6A and M10A, highlighting (positive and negative) regulation of pri-miRNA transcription by RNA polymerase II. M9A reflected viral processes (viral budding via the host endosomal sorting complexes required for transport (ESCRT) and virion assembly), whilst M8A comprised negative regulation of epithelial/endothelial cell apoptosis. M7A, the smallest cluster, was associated with on protein localization and targeting to membranes. M3A captured Hippo-pathway regulation and adenylate cyclase activation, while M5A was associated with nucleotide metabolism, particularly pyrimidine and nucleoside monophosphate phosphorylation.

**Fig. 3:**
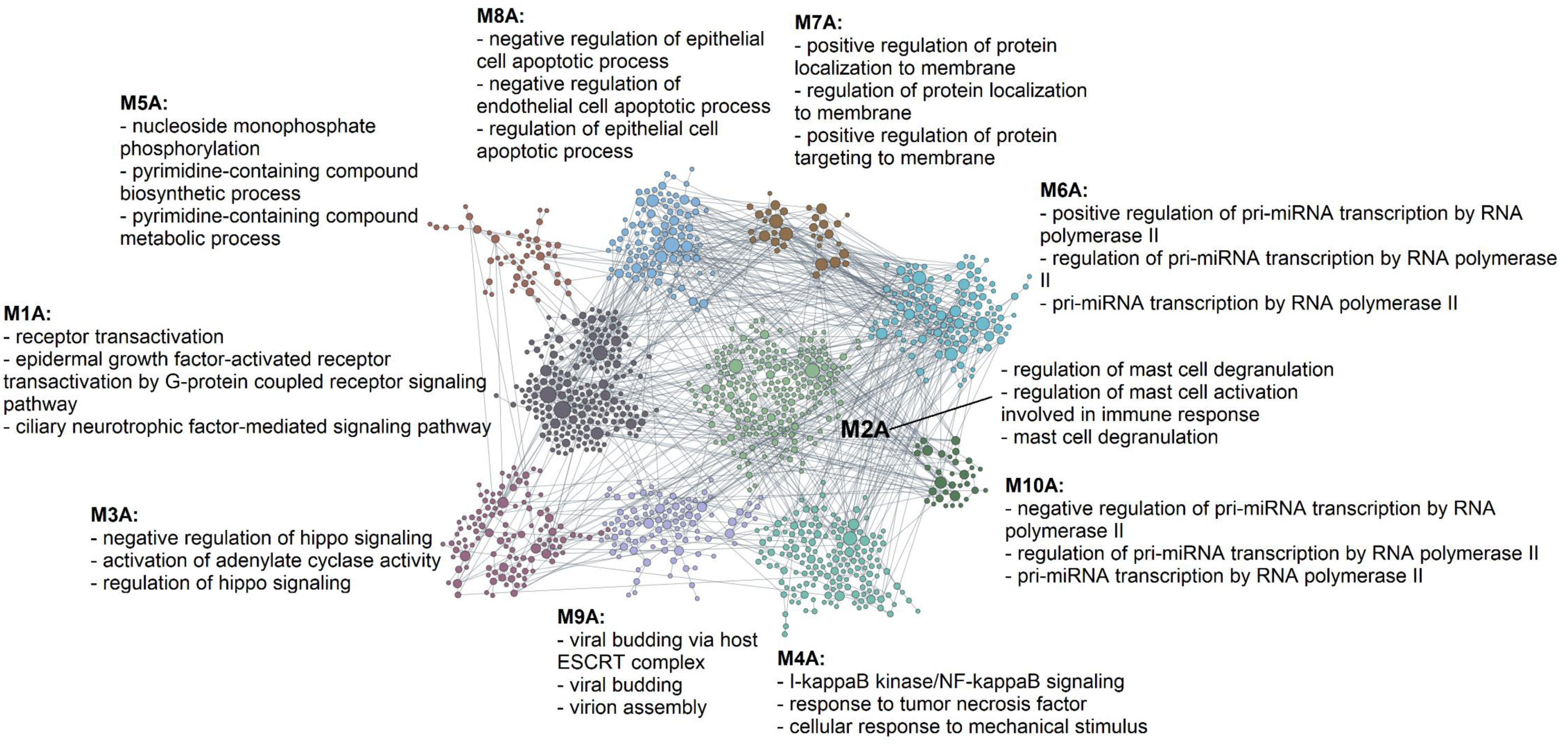
Functional sub-module discovery of the yellow module highlights ten sub-modules associated with specific pathways. Using a functional module discovery method within the HumanBase platform we identified clusters of genes within the yellow module that show co-ordinated activity in gene ontology (GO) biological processes. Nodes are genes grouped into co-functional sub-modules (labelled M1A-M10A). Labels list representative enriched GO Biological Process terms for each sub-module.

In the tan module associated with infection, our analysis resolved three sub-modules (M1B–M3B) with distinct biological processes captured (Fig. 4). M1B reflected host–pathogen transcriptional interactions, including modulation by the host of viral and symbiont transcription. M2B reflected steroid-responsive and epidermal programmes, enriched for cellular responses to glucocorticoid/corticosteroid stimuli and regulation of keratinocyte proliferation. M3B represented kinase-signalling and developmental processes, enriched for positive regulation of peptidyl-threonine phosphorylation and pituitary gland development.

**Fig. 4:**
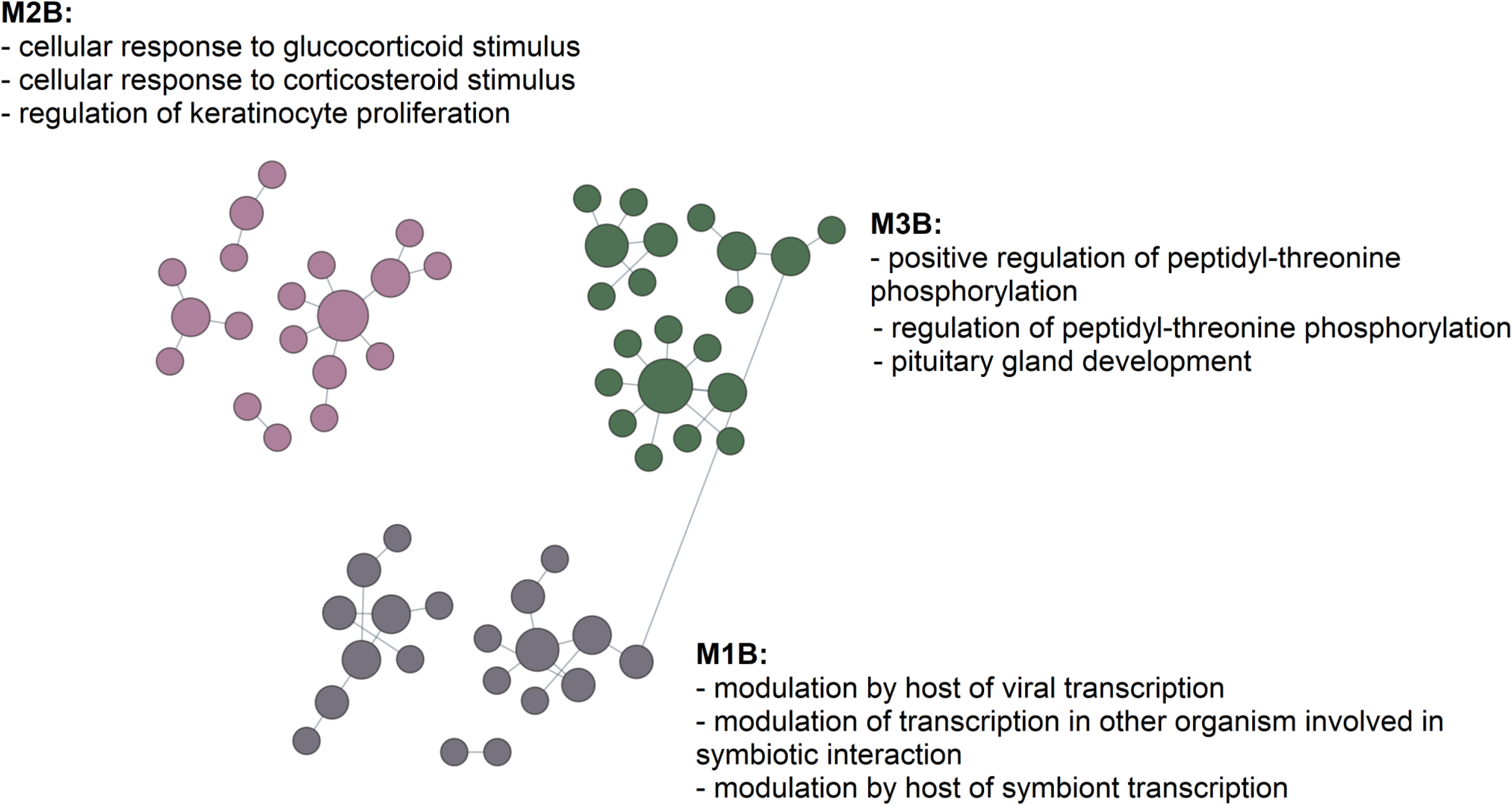
Functional sub-module discovery of the tan module highlights three sub-modules associated with specific pathways. Using a functional module discovery method within the HumanBase platform we identified clusters of genes within the tan module that show co-ordinated activity in gene ontology (GO) biological processes. Nodes are genes grouped into co-functional sub-modules (labelled M1B-M3B). Labels list representative enriched GO Biological Process terms for each sub-module.

In the red module, which had shown a Bonferroni-significant correlation with AD status (Figure 2A), we identified four sub-modules (M1C-M4C) (Fig. 5), revealing pathways and processes well recognized in AD, including regulation of Notch signalling pathways (M3C sub-module) and those associated with the mitochondria, mitosis, and ubiquitination (M4C sub-module). The other two sub-modules were largely related to cellular response to cadmium and diamide (M2C) and negative regulation of phosphatidylinositol 3-kinase (PI3K) and lipid kinase activity (M1C sub-module), which have also been associated previously with AD (Kumar and Bansal, 2021).

**Fig 5:**
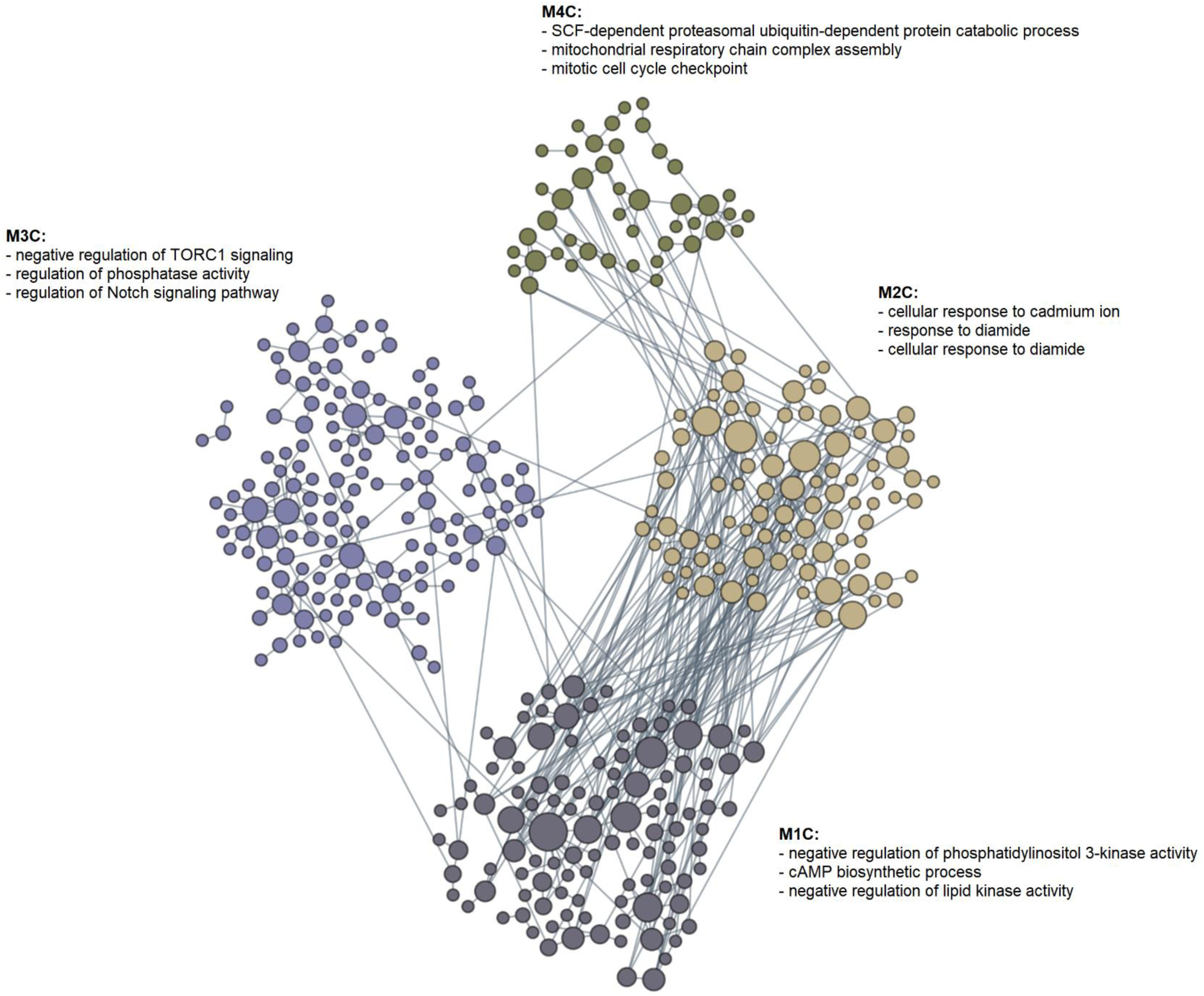
Functional sub-module discovery of the red module highlights four sub-modules associated with specific pathways. Using a functional module discovery method within the HumanBase platform we identified clusters of genes within the red module that show co-ordinated activity in gene ontology (GO) biological processes. Nodes are genes grouped into co-functional sub-modules (labelled M1C-M4C). Labels list representative enriched GO Biological Process terms for each sub-module.

## Discussion

In recent years, a growing number of studies have explored cortical transcriptomic signatures in AD compared to control donors (Mathys et al., 2019; Mostafavi et al., 2018; Q et al., 2021; Raj et al., 2018; Wan et al., 2020) however, these studies have not explored whether concurrent systemic infection impacts these profiles, which is important given the clinical links between systemic infections and AD manifestation. First, we set out to explore transcriptional signatures in AD compared to control prefrontal cortex in the absence of systemic infection, identifying 763 significant DEGs, which were enriched in bioenergetic and oxidative phosphorylation pathways. These pathways have already been reported as dysregulated in neurodegenerative diseases, including AD (Manczak et al., 2004; Shoffner, 1997). KEGG enrichment further highlighted an FDR significant enrichment in pathways implicated in HD and prion disease. This likely represents an enrichment of neurodegenerative disease processes, as the majority of genes in these pathways overlapped with each other and also with the KEGG AD pathway, which was nominally significantly enriched. Together, these findings indicate that our transcriptomic analysis of AD and control brains in the absence of systemic infection captured core neurodegenerative processes characteristic of AD pathology. Interestingly, the most significant gene we identified was *ADAMTS2*, which has been consistently reported in transcriptomic studies of AD (McCorkindale et al., 2022; Q et al., 2021; Tsatsanis et al., 2021). Several other highly ranked genes also overlap with previous AD findings, including *PPDPF* (McCorkindale et al., 2022; Mitra et al., 2024; Q et al., 2021), *HEY1* (Leal et al., 2012) and *VGF* (Hölttä et al., 2015; Yu et al., 2023).

Our study is the first to stratify analyses by the presence of a systemic respiratory infection at the time of death. We identified 122 significant DEGs between AD and control cortex in the presence of a systemic infection. The most significant gene was *LPIN2,* which has been previously reported to be downregulated in COVID-19 patients and functions as a regulator of inflammation during viral infection, attenuating toll-like receptor 3 (TLR3) signalling, reactive oxygen species (ROS) production, and mitochondrial DNA release, thereby limiting activation of the NLRP3 inflammasome (de Pablo et al., 2023).

Interestingly, we identified 57 genes that were significantly differentially expressed only in AD compared to controls with infection and were not altered in uninfected cases. Among these, 18 showed no nominally significant transcriptional change in the absence of infection (p > 0.05). One of these genes is *MAPK4* (mitogen-activated protein kinase 4), which has been implicated in host– pathogen interactions and host-cell survival. *MAPK4*-deficient human cells exhibit reduced parasite infection (Watanabe et al., 2023). Another, *VAV3,* is a regulator of endothelial barrier integrity, including in brain microvascular endothelial cells (Hilfenhaus et al., 2018), whilst *POU3F4* has been reported to be upregulated in human cells following *R. prowazekii* infection (Ge et al., 2008).

Beyond individual gene-level changes, GSVA highlighted distinct shifts in biological pathway activity that further illuminate the impact of systemic infection in AD brain. GSVA revealed distinct pathway activity between AD and controls, depending on the infection status. In AD without infection, pathways such as “L-arginine transmembrane transport”, “regulation of smoothened signalling”, “embryonic organ development”, and “B-cell receptor signalling pathway” were upregulated, suggesting the reactivation of developmental and immune programs. At the same time, AD brains showed downregulation of “cortical cytoskeleton organization”, “cellular response to estrogen stimulus”, and “antimicrobial humoral immune response mediated by antimicrobial peptide”, pointing to impaired cellular homeostasis and weakened host defences. During systemic infection, the AD transcriptome shifted, with upregulation of “cell development”, “regulation of DNA-binding transcription factor activity”, “positive regulation of lipid biosynthetic process”, “DNA-templated transcription”, and “regulation of signal transduction by p53 class mediator”, potentially reflecting an enhanced stress and repair response. In parallel, infection in AD was linked to suppression of antimicrobial humoral pathways and circulatory system development, indicating dampened immune response and vascular development functions. Together, these results suggest that infection does not simply amplify existing AD signatures but instead potentially redirects transcriptional changes.

Examining gene networks provides additional context for understanding the effects of systemic infection in AD. WGCNA identified three important modules: the red module, which showed a Bonferroni-significant association with AD, the yellow module, nominally associated with both AD and infection, and the tan module, nominally associated with infection. The red module showed an enrichment in KEGG pathways related to aging, signal transduction and endocrine systems. Notably, when we performed functional sub-module analysis, we identified sub-modules related to pathways previously associated with AD, including Notch signalling and those associated with the mitochondria, mitosis and ubiquitination.

GO enrichment analysis of the yellow module identified pathways such as “small GTPase– mediated signal transduction”, “actin filament organisation”, and “establishment of cell polarity”, pointing to cytoskeletal remodelling and cell signalling. These processes are fundamental for maintaining BBB structure and function. Additionally, pathogens have been reported to promote reorganization of the cytoskeleton through the activity of GTPases to enter the host cells. In contrast, GO enrichment analysis of the tan module highlighted numerous pathways including those related to glia, stress responses and immune regulation, like “cellular response to immune stimulus”, “cellular response to platelet-derived growth factor stimulus”, “glial cell proliferation” and “regulation of reactive oxygen species metabolic processes”. Unlike the yellow module, which correlates with both AD and infection, the tan module was only associated with infection, suggesting that its signature reflects infection-driven changes rather than changes in the context of AD and infection. The enrichment in astrocytes, pericytes, and endothelial cells further points to infection-related alterations in the BBB, where these cell types jointly regulate vascular integrity and immune cell trafficking.

Focusing on the yellow module, the functional sub-module analysis revealed ten modules grouped by co-functional gene sets. These submodules capture diverse biological processes, ranging from receptor transactivation and EGFR–GPCR signalling to Hippo pathway regulation, NF-κB–mediated inflammatory signalling, viral budding via the ESCRT complex, pri-miRNA transcriptional regulation, apoptosis and protein localization to the membrane. This diversity highlights that the yellow module is not driven by a single pathway but rather reflects an integrated network of cytoskeletal remodelling, transcriptional regulation, and host–pathogen interaction processes. This overlaps with pathways shown to have altered activity in our GSVA analysis of AD compared to controls during infection, where we also saw alterations in cytoskeleton and signalling pathways. The tan module, which was specifically associated with infection but not with AD, contained submodules related to host modulation of viral transcription and glucocorticoid/stress responses, again overlapping with our GSVA pathway activity analysis, which had shown alterations in immune and vascular stress responses in AD compared to control during infection.

While these insights advance our understanding of not only AD but also the brain’s response to systemic infection, some limitations should be noted. First, there are limitations related to the use of post-mortem brain tissue, as this is a cross-sectional analysis and does not allow us to explore the cumulative effect of infections over the life course. As a result, we were unable to explore whether recent (but resolved) systemic infections may result in brain transcriptional alterations. Second, despite efforts to exclude comorbidities, a small number of AD cases also showed evidence of vascular pathology from the post-mortem records. Third, due to sample availability there was a small imbalance in sample sizes between groups, with a smaller number of control donors with a systemic infection (N = 37) compared to controls without infection (N = 52). This may have resulted in reduced power to detect changes in our differential gene expression analysis comparing AD and control donors with an infection and may be the reason we identified fewer FDR-significant genes (N = 122) in this analysis. It is also worth noting that although we were able to identify a Bonferroni-significant WGCNA module associated with AD, the modules that were correlated with infection did not pass multiple testing correction. Finally, the use of bulk brain tissue is also a caveat of our study. Although we controlled for cell proportions in our analyses and performed cell type enrichment analysis for modules of interest, future studies should be undertaken on sorted cell populations or at the level of the single-cell to reduce heterogeneity. In addition, future directions should explore whether different types of infection are associated with activation of specific molecular pathways in the brain. Ultimately, studies incorporating multi-omics approaches, integrating genetic, epigenetic proteomic data from the same samples will be important, will allow a holistic assessment of the impact of systemic infections on the brain in AD.

In conclusion, this study emphasises the intricate molecular and cellular interactions occurring in the brain during AD and systemic infection. These findings broaden the microglia-centric perspective of neuroinflammation in AD, highlighting that systemic infections in AD, may alter vascular and immune pathways and engage host-pathogen processes in BBB-associated cell types, particularly astrocytes and pericytes. This is particularly important given the clinical links between systemic infections and worsened cognition and outcomes in AD. This study provides a good foundation for future research exploring the potential of therapeutic strategies that mitigate neuroinflammation and repair vascular dysfunction in AD.

## Author Contributions

**Giulia Pegoraro**: Methodology, Software, Formal Analysis, Data Curation, Writing – Original Draft, Vizualization; **Lachlan F. MacBean**: Investigation; **Adam R. Smith**: Investigation, Data Curation, Writing – Review & Editing, Project administration; **Darren M. Soanes**: Software, Formal Analysis; **Rebecca G. Smith**: Data Curation, Funding acquisition; **Jennifer Imm**: Investigation; **Joshua Harvey**: Investigation, Software; **Morteza Kouhsar**: Software; **Valentin Laroche**: Software, Formal Analysis, **Luke Weymouth:** Investigation, Data Curation; **Isabel Castanho**: Supervision; **Winston Hide**: Supervision; **Laura Palmer**: Resources; **Debra J. Lett**: Resources; **Andrew C. Robinson**: Resources; **Natalie Woodman**: Resources; **Karina McDade**: Resources; **Delphine Boche**: Conceptualization, Funding acquisition; **Ehsan Pishva**: Supervision, Writing – Review & Editing; **Katie Lunnon**: Conceptualization, Writing – Original Draft, Supervision, Project administration, Funding acquisition.

---

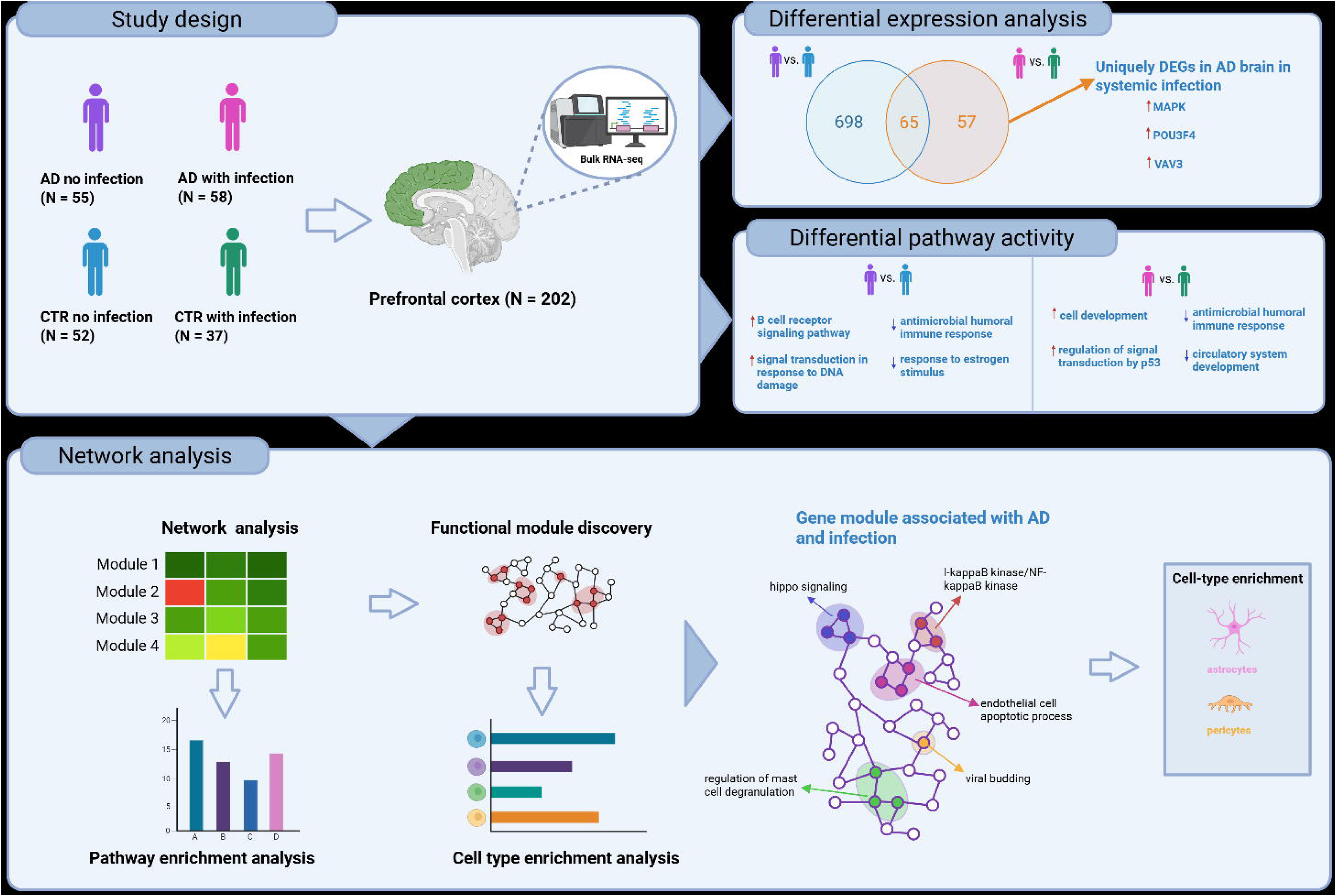

